# HiCcompare: a method for joint normalization of Hi-C datasets and differential chromatin interaction detection

**DOI:** 10.1101/147850

**Authors:** John C. Stansfield, Mikhail G. Dozmorov

**Author notes:** Corresponding author (MGD).

## Abstract

Changes in spatial chromatin interactions are now emerging as a unifying mechanism orchestrating regulation of gene expression. Evolution of chromatin conformation capture methods into Hi-C sequencing technology now allows an insight into chromatin interactions on a genome-wide scale. However, Hi-C data contains many DNA sequence- and technology-driven biases. These biases prevent effective comparison of chromatin interactions aimed at identifying genomic regions differentially interacting between, disease-normal states or different cell types. Several methods have been developed for normalizing individual Hi-C datasets. However, they fail to account for biases *between two or more Hi-C datasets*, hindering comparative analysis of chromatin interactions. We developed a simple and effective method `HiCcompare` for the joint normalization and differential analysis of multiple Hi-C datasets. The method avoids constraining Hi-C data within a rigid statistical model, allowing a data-driven normalization of biases using locally weighted linear regression (loess). The method identifies region-specific chromatin interaction changes complementary to changes due to large-scale genomic rearrangements, such as copy number variants (CNVs). `HiCcompare` outperforms methods for normalizing individual Hi-C datasets in detecting *a priori* known chromatin interaction differences in simulated and real-life settings while detecting biologically relevant changes. `HiCcompare` is freely available as a Bioconductor R package https://bioconductor.org/packages/HiCcompare/.

**Author Summary:** Advances in chromosome conformation capture sequencing technologies (Hi-C) have sparked interest in studying the 3-dimensional (3D) chromatin interaction structure of the human genome. The 3D structure of the genome is now considered as a primary regulator of gene expression. Changes to the 3D chromatin interactions are now emerging as a hallmark of cancer and other complex diseases. With the growing availability of Hi-C data generated under different conditions (e.g. tumor-normal, cell-type-specific), methods are needed to compare them. However, biases in Hi-C data hinder their comparative analysis. To account for biases, several normalization techniques have been developed for removing biases in individual Hi-C datasets, but very few were designed to account for between-datasets biases. We developed a new method and R package `HiCcompare` for the joint normalization of multiple Hi-C datasets and differential chromatin interaction detection. Our results show the superiority of our joint normalization methods compared to methods for normalizing individual datasets in detecting true chromatin interaction changes. `HiCcompare` enables further research into discovering the dynamics of 3D genomic changes.

## Introduction

The 3D chromatin structure of the genome is emerging as a unifying regulatory framework orchestrating gene expression by bringing transcription factors, enhancers and co-activators in spatial proximity to the promoters of genes [1–10]. Together with epigenomic profiles, changes in chromatin interactions shape cell type-specific gene expression [11–17], as well as misregulation of oncogenes and tumor suppressors in cancer [1,18–20]. Identifying changes in chromatin interactions is the next logical step in our understanding of genome regulation.

Development of Chromatin Conformation Capture (3C) sequencing technology [21] and its derivatives continues to help better understand the 3D structure of the genome [14]. As such technologies require significant labor, sequencing, and data storage costs, a variety of simplified technologies and corresponding statistical methods for data analysis have been developed (e.g., ChIA-PET [19], Capture Hi-C [22]). Although valuable for understanding long-distance interactions (e.g., promoters vs. enhancers), these technologies provide only a partial view of the 3D structure. In contrast, Hi-C technology allows the detection of “all vs. all” long-distance chromatin interactions across the whole genome [14,23,24]. It proved to be indispensable for advancing our understanding of copy number variations, long-range epigenetic remodeling, and atypical gene expression corresponding to disrupted chromatin interactions in cancer [18,25]. Hi-C technology is becoming the flagship approach for understanding the 3D organization of the genome, signifying the need for proper computational methods for its analysis [26].

Soon after public Hi-C datasets became available, it was clear that technology- and DNA sequence-driven biases substantially affect chromatin interactions [27]. The technology-specific biases include cutting length of a restriction enzyme (HindIII, MboI, or NcoI), cross-linking conditions, circularization length, etc. [28,29]. The DNA sequence-driven biases include GC content, mappability, nucleotide composition [27]. Discovery of these biases led to the development of methods for normalizing individual datasets [14,27,30,31]. Although normalization of individual datasets improves reproducibility within replicates of Hi-C data [27,30,32], these methods do not consider biases between multiple Hi-C data.

Accounting for the between-datasets biases is critical for the correct identification of chromatin interaction changes between, e.g., disease-normal states, or cell types. Left unchecked, biases can be mistaken for biologically relevant differential interactions. While DNA sequence-driven biases affect the data similarly (e.g., CG content of genomic regions tested for interaction differences is the same), technology-driven biases are poorly characterized and affect chromatin interactions unpredictably. Importantly, another source of chromatin interaction differences is due to large-scale genomic rearrangements, such as copy number variations [33,34], a frequent event in cancer genomes [35,36]. Accounting for such biases is needed for the detection of differential chromatin interactions between Hi-C datasets.

We developed an R package `HiCcompare` for the joint normalization and comparative analysis of multiple Hi-C datasets, summarized as chromatin interaction matrices. Our method is based on the observation that chromatin interactions are highly stable [37–40], suggesting that the majority of them, excluding large genomic rearrangement of regions, can serve as a reference to build a rescaling model. We present the novel concept of the MD plot (difference vs. distance plot), a modification of the MA plot [41], visualizing the distance-centric differences between interacting chromatin regions, where distance is expressed in terms of unit-length size of the regions. The MD plot concept naturally allows for fitting the local regression model, a procedure termed `loess`, and jointly normalizing the two datasets by balancing biases between them. The distance-centric view of chromatin interaction differences allows for detecting statistically significant differential chromatin interactions between two Hi-C datasets using a simple but robust permutation method. We show improved performance of differential chromatin interaction detection when using the jointly vs. individually normalized simulated and real Hi-C datasets. Our method is broadly applicable to a range of biological problems, such as identifying differential chromatin interactions between tumor and normal cells, immune cell types, and normal tissues.

## Results

### The off-diagonal concept of distance between regions in chromatin interaction matrices

Our study focuses on the analysis of multiple Hi-C datasets by providing functions for the joint normalization of two or more chromatin interaction matrices and differential chromatin detection among them. We provide a method for analyzing processed Hi-C sequencing data, i.e., summarized as two-dimensional matrices of chromatin interaction frequencies. Briefly, Hi-C processing pipelines summarize the data into matrices by binning each chromosome into discrete regions - ‘windows’ - that define the resolution (unit-length) of the data. Each row and column in a chromatin interaction matrix corresponds to a region. Each cell contains a number of reads shared between each pair of genomic regions, a proxy for interaction frequency. Chromosome-specific interaction matrices are symmetric; inter-chromosomal matrices are oblong due to the differing number of ‘windows’ on different chromosomes. The frequency of inter-chromosomal interactions is much smaller and much less consistent [22,42,43]. Consequently, inter-chromosomal interaction matrices contain a large proportion of zeros. In this study, we focus on normalization and differential analysis of chromosome-specific interaction matrices. However, the concept is applicable to inter-chromosomal normalization/comparison and will be extended to the analysis of the whole-genome chromatin interaction matrices.

A foundation of our methods is the distance-centric view of chromosomal interactions. Fig 1A illustrates the concept of the unit-length distance between interacting regions captured by the adjacency matrix. The values on the diagonal trace represent interaction frequencies of self-interacting regions. Each off-diagonal set of values represents interaction frequencies for a pair of regions at a given unit-length distance. The unit-length distance is expressed in terms of resolution of the data (the size of interacting regions). For data at 10kb resolution, the first off-diagonal set of values represents chromatin interaction frequencies between regions spaced at 10kb, etc. Regions closer to each other in a linear space tend to interact more frequently [14,22], as illustrated by the color intensity near the diagonal trace. The average interaction frequency, measured at each off-diagonal unit-length distance, drops as the distance between interacting regions increases. The concept of considering interaction frequencies at each distance, represented by values at each off-diagonal trace in chromatin interaction matrices, is central for the joint normalization and differential chromatin interaction detection.

**Fig 1.**
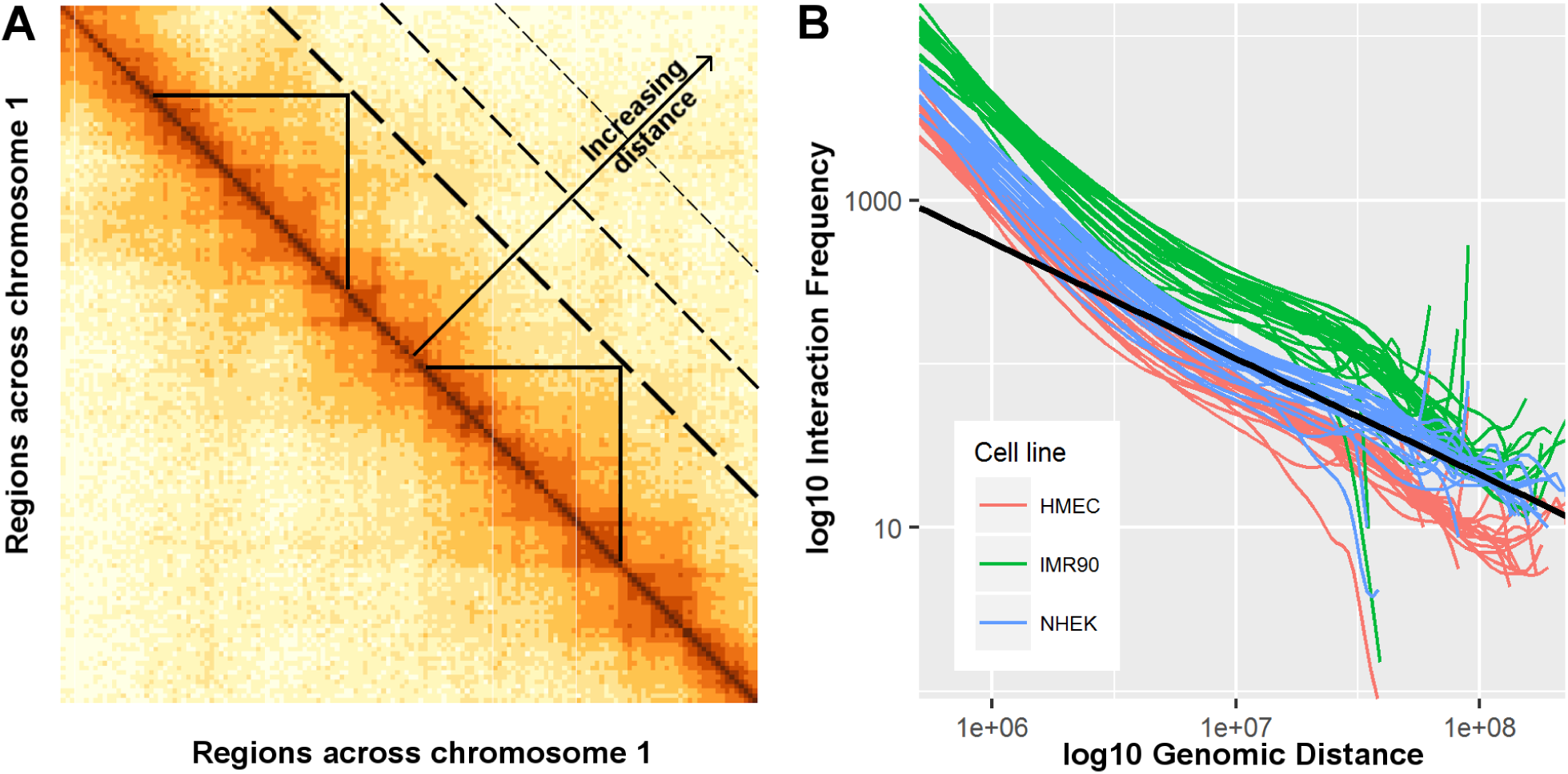
Chromatin interaction frequency representation and dependence on distance. (A) Distance-centric (off-diagonal) view of chromatin interaction matrices. Values on the diagonal represent interaction frequencies of regions at distance 0 (self-interacting). Each off-diagonal vector of interaction frequencies represents interactions at a given distance between pairs of regions. Triangles mark pairs of genomic regions interacting at the same distance. Data for chromosome 1, K562 cell line, 50KB resolution, spanning 0 - 7.5Mb is shown. (B) Deviation from the ideal power-law relationship (straight lines) between the *log*_10_*-log*_10_ interaction frequencies and distance. Curved lines represent chromosome-specific `loess` fits of the relationship, colored by datasets. The full range of genomic distances is shown. Data from HMEC, IMR90, NHEK cell lines, using all chromosomes, 500kb resolution were used.

### Non-parametric relationship between chromatin interaction frequencies and distance

Numerous attempts have been made to parametrically model the inverse relationship between chromatin interaction frequency and the distance between interacting regions. These include power-law [14,24], double exponential [44], binomial [45], Poisson and negative binomial [9,13,22,32], and zero-inflated negative binomial [46] distributions. These distributions are then used to identify regions that interact significantly stronger than would be predicted by the model [9,13,22,32,32,45].

The aforementioned publications acknowledge that parametric approaches fail to model chromatin interaction frequencies across the full range of distances between interacting regions, even within the same chromosome [24]. Our analysis of real Hi-C data confirms that each chromosome has a unique distribution of chromatin interaction frequencies vs. distances (Fig 1B, S1 Fig), complicating the use of a single model to approximate interaction frequency vs. distance dependence. Consequently, the parametric assumptions used to normalize individual Hi-C datasets may be violated, justifying the need for the non-parametric approach for normalization of Hi-C data.

### Persistence of biases in individually normalized Hi-C replicated data

Methods for normalizing individual Hi-C datasets can be broadly divided into two categories. The first category relies on parametric modeling of chromatin interaction frequencies that helps to adjust for biases deviating from the model. The second, matrix-balancing techniques, assume the cumulative effect of bias is captured in the global chromatin interaction matrix. Both categories aim to alleviate biases in individual chromatin interaction matrices by uniformly adjusting them. However, when comparing two or more chromatin interaction matrices, it is unclear whether methods that normalize individual datasets can eliminate biases *between* the datasets.

Biases due to large scale genomic variations, such as copy number variants (CNVs), introduce another layer of complexity only recently recognized in normalization methods of individual Hi-C datasets [34]. Such genomic events will introduce *en masse* chromatin interaction changes, overshadowing the region-specific changes in chromatin interactions. Our framework has been designed to detect region-specific chromatin interaction differences; therefore, genomic regions containing CNVs and other large-scale changes should be detected separately [33] and analyzed in parallel.

To assess the between-datasets biases, we introduce the novel concept of an MD plot (see Methods), designed to visualize two Hi-C datasets on one plot. Briefly, differences in chromatin interaction frequencies (**M**inus) are visualized on a per-unit-length distance basis (Fig 2A). Owing to the fact that chromatin interactions are highly conserved [37–39], we expect that the majority of the **M** differences should be relatively unchanged among the Hi-C datasets (centered around **M** equal to zero). The MD plot visualization allows us to identify systematic biases appearing as the offset of the cloud of **M** differences from zero. Visualizing replicates of Hi-C data (Gm12878 cell line) showed the presence of biases (Fig 2A). Importantly, these biases persisted in the individually normalized datasets (Fig 2C-F), suggesting that the performance of methods normalizing individual matrices may be sub-optimal when comparing multiple Hi-C datasets.

**Fig 2.**
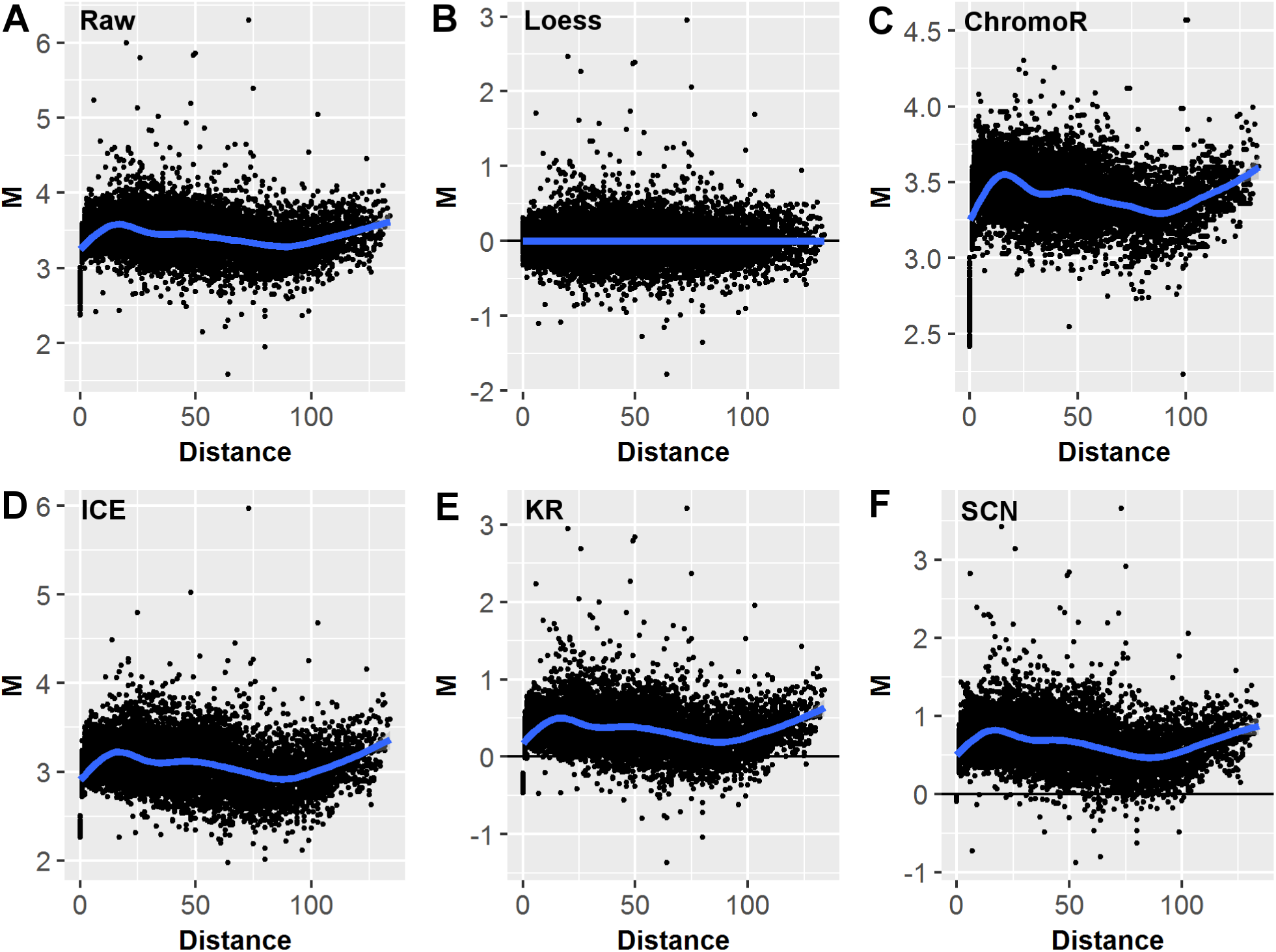
MD plot data visualization and the effects of different normalization techniques. MD plots of the differences *M* between two replicated Hi-C datasets (GM12878 cell line, chromosome 11, 1MB resolution, DpnII and MboI restriction enzymes) plotted vs. distance *D* between interacting regions. (A) Before normalization, (B) after `loess` joint normalization, (C) `ChromoR`, (D) Iterative Correction and Eigenvector decomposition (`ICE`), (E) Knight-Ruiz (`KR`), (F) Sequential Component Normalization (`SCN`).

### Elimination of biases in jointly normalized Hi-C datasets

To account for the between-datasets biases, we developed a non-parametric joint normalization method that makes no assumptions about the theoretical distribution of the chromatin interaction frequencies. It utilizes the well-known `loess` (locally weighted polynomial regression) smoothing algorithm - a regression-based method for fitting simple models to segments of data [47]. `loess` has a well-established history in microarray data analysis, normalizing red-green gene expression channels, or adjusting gene expression between pairs of single-channel arrays [48]. With the advent of sequencing technologies, `loess` has been applied to normalize pairs of ChIP-seq datasets [49,50] and Hi-C data [51]. The main advantage of `loess` is that it accounts for any local irregularities *between* the datasets that cannot be modeled by parametric methods. Thus, `loess` is particularly appealing when normalizing two Hi-C datasets, as the internal biases in Hi-C data are poorly understood.

In contrast to parametric methods, `loess` makes no *a priori* assumptions about chromatin interaction frequencies, thus allowing the Hi-C data to self-guide the normalization process (see Methods). Using the data from an MD plot, it minimizes the systematic biases between the datasets, while preserving local and, potentially, biologically significant chromatin interaction differences. Applied to real Hi-C replicated data, it successfully eliminated biases (Fig 2B). On the contrary, biases remained in the individually normalized datasets, hindering their comparison and the detection of differentially interacting chromatin regions (S1 File).

Importantly, the between-dataset chromatin interaction changes may be due to large-scale genomic rearrangements and copy number variants (CNVs), a frequent case in tumor-normal comparisons [36,51]. The effect of CNVs on Hi-C data normalization has been recognized [34]. Therefore, we recommend excluding genomic regions containing CNVs from the joint normalization, so only copy-number neutral genomic regions are used. This strategy will lead to optimal performance of the joint normalization method, while allowing for CNV-driven changes to be analyzed in parallel.

### Per-unit-length-distance concept of detecting differential chromatin interactions using permutation framework

To the best of our knowledge, only three methods attempted the comparative analysis of Hi-C data. The `diffHic` method [51] is an extension of the popular RNA-seq differential expression method `edgeR`, operating on raw sequencing data. As such, it leaves the user with challenges of sequencing data storage, processing, normalization, summarization, and other bioinformatics heavy lifting of Hi-C data. The `HiCCUPS` algorithm [39] searches for clusters of chromatin interaction “hotspots” in individual matrices - entries in which the frequency of contacts is enriched relative to the local background. The “hotspots” different between two chromatin interaction matrices are identified by intersection, which does not address the significance of the differences and leaves the problem of between-datasets biases unaddressed. The only method to statistically compare processed Hi-C dataset is `ChromoR` [9]. However, in our tests it has failed to detect differential chromatin interactions in real Hi-C data, perhaps due to the use of the parametrically constrained model (S1 File), an approach that has been criticized [52]. The lack of methods for detecting statistically significant chromatin interaction differences between processed Hi-C matrices prompted us to develop a new simple differential chromatin interaction detection algorithm.

To detect significant chromatin interaction differences, we used the representation of the differences in the MD coordinate system. Importantly, the MD plot naturally prompts testing of the differences on a per-unit-length-distance basis, an idea we incorporated into a per-unit-distance permutation framework (see Methods). Briefly, distance *d*-specific vectors of chromatin interaction differences *M*_*d*_ are used to provide a reference distribution to calculate the probability of detecting a given difference, or larger. The permutation framework naturally accounts for multiple testing. Such a simple approach showed excellent performance in detecting *a priori* introduced chromatin interaction differences, even when the data is normalized using individual normalization methods (S5-S6 Files).

### loess joint normalization improves differential chromatin interaction detection

The power of differential chromatin interaction detection was first assessed using simulated Hi-C matrices with controlled chromatin interaction differences. We simulated pairs of chromatin interaction maps that contain interaction frequencies and biases resembling the real data (see Methods and S2-S4 Files), and introduced controlled fold changes in one of them. The matrices were normalized using the `loess` and `MA` joint normalization methods, and each of the four methods (`ChromoR`, `KR`, `ICE`, `SCN`, see Methods for the brief description of each) for normalizing individual matrices.

The ROC curve analysis showed the superiority of the `loess` joint normalization in improving the power of detecting differential chromatin interactions across the range of controlled fold changes (Fig 3). The benefits of the `loess` joint normalization were the most pronounced at detecting lower fold changes (Fig 3A). Notably, the `MA` normalization was second in performance to the `loess` joint normalization, strengthening the need to normalize Hi-C datasets jointly. Interestingly, the performance of the non-normalized data was equal to or better than all normalization methods except the `loess` joint normalization, questioning the need for normalization for the differential chromatin interaction detection. Expectedly, higher fold changes ≥4 were easier to detect, as reflected by the relatively good performance of all but `ChromoR` normalization methods. The benefits of normalization as compared with the non-normalized data were easier to detect at the higher fold changes, with the `loess` joint normalization performing best (Fig 3B-D). Among methods for normalization of individual chromatin interaction matrices the `KR` method performed best, following the `ICE` and `SCN` methods (Fig 3). Surprisingly, the `ChromoR` method performed the worst, confirming our observation of its poor performance in removing biases and detecting differential chromatin interactions when used alone (S1 File).

**Fig 3.**
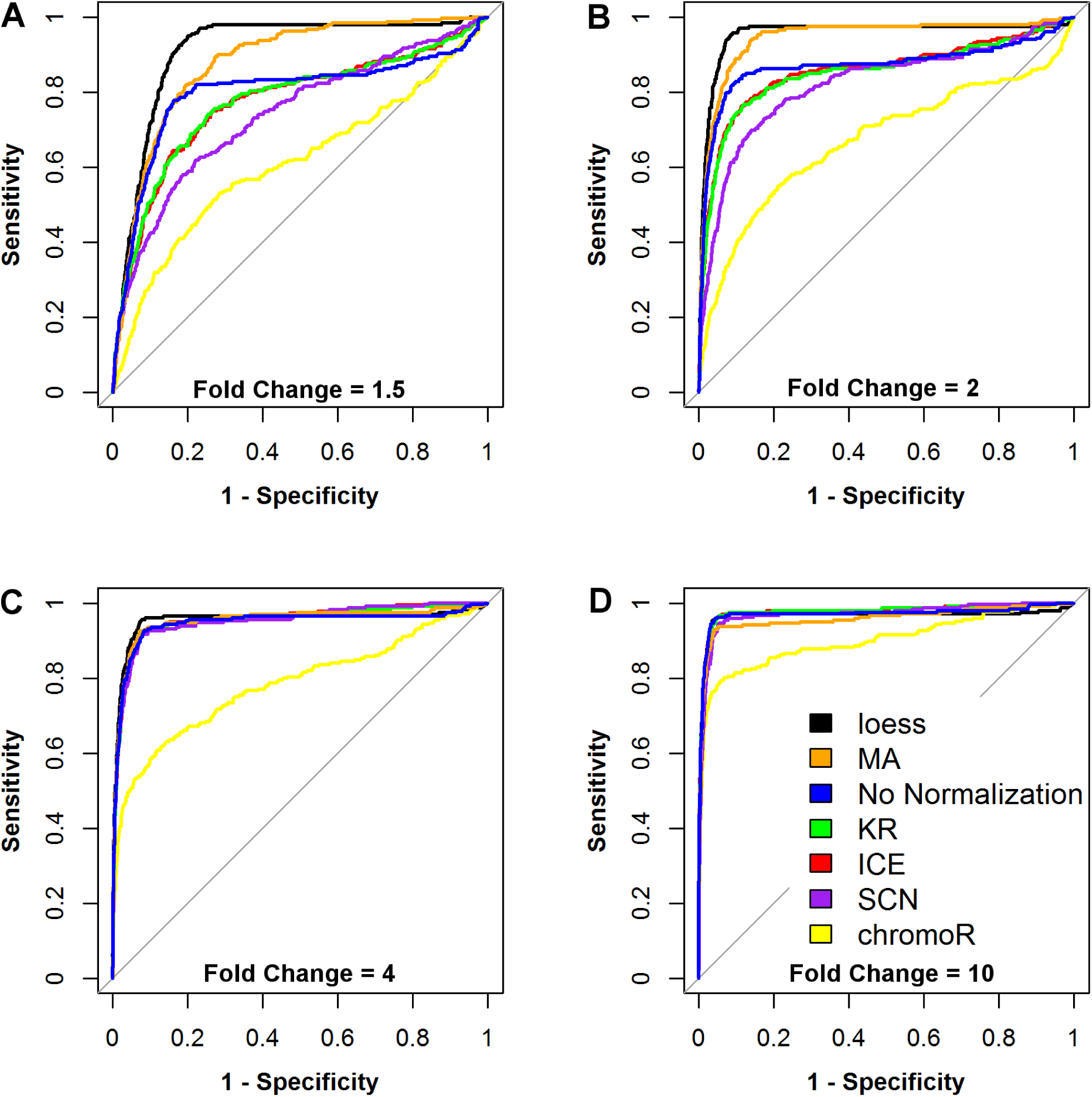
ROC curves for different normalization techniques. ROC curves of the differential chromatin interaction detection using different normalization techniques at (A) 1.5, (B) 2.0, (C) 4.0, (D) 10.0 fold changes. Simulated 100 x 100 chromatin interaction matrices with 250 controlled changes were used.

As no single metric can evaluate all aspects of classifier performance [53], we evaluated the performance of the normalization methods using additional metrics. Confirming our observation that the joint normalization method yields the largest area under the curve (Fig 3), it also had the highest true positive rate (TPR), the smallest false discovery rate (FDR), improved accuracy and precision, as compared with methods for normalizing individual Hi-C matrices (S5 File). In summary, the `loess` joint normalization outperformed individual normalization methods in improving the power of detecting differential chromatin interactions.

Controlled changes may also be introduced into replicates of real Hi-C data. Replicated experiments are assumed to contain minimal differences, primarily due to technical noise. However, replicates of real data contain many more differentially interacting chromatin regions (∼45%, [39]) than simulated data due to the much larger effect of biases. These existing differences may often be larger than the controlled fold changes and therefore detected as false positives if biases remain unaccounted for. Indeed, all but `loess` normalization methods performed sub-optimally when controlled changes less than 2-fold were introduced in the replicates of real Hi-C data (S2 Fig). Similar to the simulated settings, the non-normalized data provided sufficient power to detect the controlled changes. While larger controlled changes can be detected when using different normalization methods, `loess` remained the most powerful in removing biases and the detection of small and large fold changes.

To account for the presence of the existing differences in the real Hi-C data we evaluated all major classification metrics [53], each providing a unique perspective on the performance of differential chromatin interaction detection. Confirming the results of the power analysis in simulated settings, matrices of real Hi-C data normalized using the `loess` joint normalization had the largest number of true positives (TP) and the highest true positive rate (TPR) when detecting 1.5 fold change, closely followed by the `KR` and `SCN` normalization at higher fold changes (S6 File). Consequently, the number of false positives (FP) and false negatives (FN), the false positive (FPR) and false negative (FNR) rates were the lowest in the matrices normalized with the `loess` joint normalization across the range of fold changes. Both accuracy and precision were the highest in the matrices normalized with the `loess` joint normalization. Similarly to the results in simulated settings, the individual normalization methods `KR` and `SCN` were the second and third normalization methods following the best performing `loess` joint normalization. In summary, the `loess` joint normalization improved the power of differential chromatin interaction detection across the whole range of fold changes, as compared with individual normalization methods.

### Comparison with diffHic

The `diffHiC` pipeline was designed to process raw Hi-C sequencing data and detect chromatin interaction differences using the generalized linear framework developed in the edgeR package [51,54]. Using Hi-C data analyzed in the `diffHiC` paper (human prostate epithelial cells RWPE1 overexpressing the EGR protein or GFP), we compared chromatin interaction differences detected by `diffHiC` with those detected by `HiCcompare`.

HiCcompare detected a total of 6,276 significantly different interactions in the RWPE1 data, excluding regions overlapping CNVs and/or blacklisted regions. `diffHic` detected a total of 5,737 significant differences. Surprisingly, although `diffHic` used CNV correction in their analysis, 2,567 (44.7%) of the detected differentially interacting regions overlapped with CNV regions detected in our analysis, and/or blacklisted regions. `diffHic` tended to detect differentially interacting regions with smaller fold changes as compared to `HiCcompare` (Fig 4, S4 Fig A, S2 Table). In contrast, `HiCcompare` focuses on detecting regions with high chromatin interaction differences that may be easier to detect and reproduce in the subsequent studies. Furthermore, `diffHic` seem to detect differential chromatin interactions at shorter distances between interacting regions, while `HiCcompare` detects differences across the full range of distances (Fig 4). Notably, when visualized in MA coordinates, the distribution of `diffHic`-detected changes resembled the distribution of `HiCcompare`-detected changes visualised in MD coordinates (S4 Fig B). These results suggest that detecting chromatin interaction differences represented in MD coordinates, as implemented in `HiCcompare`, may be useful in detecting large chromatin interaction differences that have more significant biological effect.

**Fig 4.**
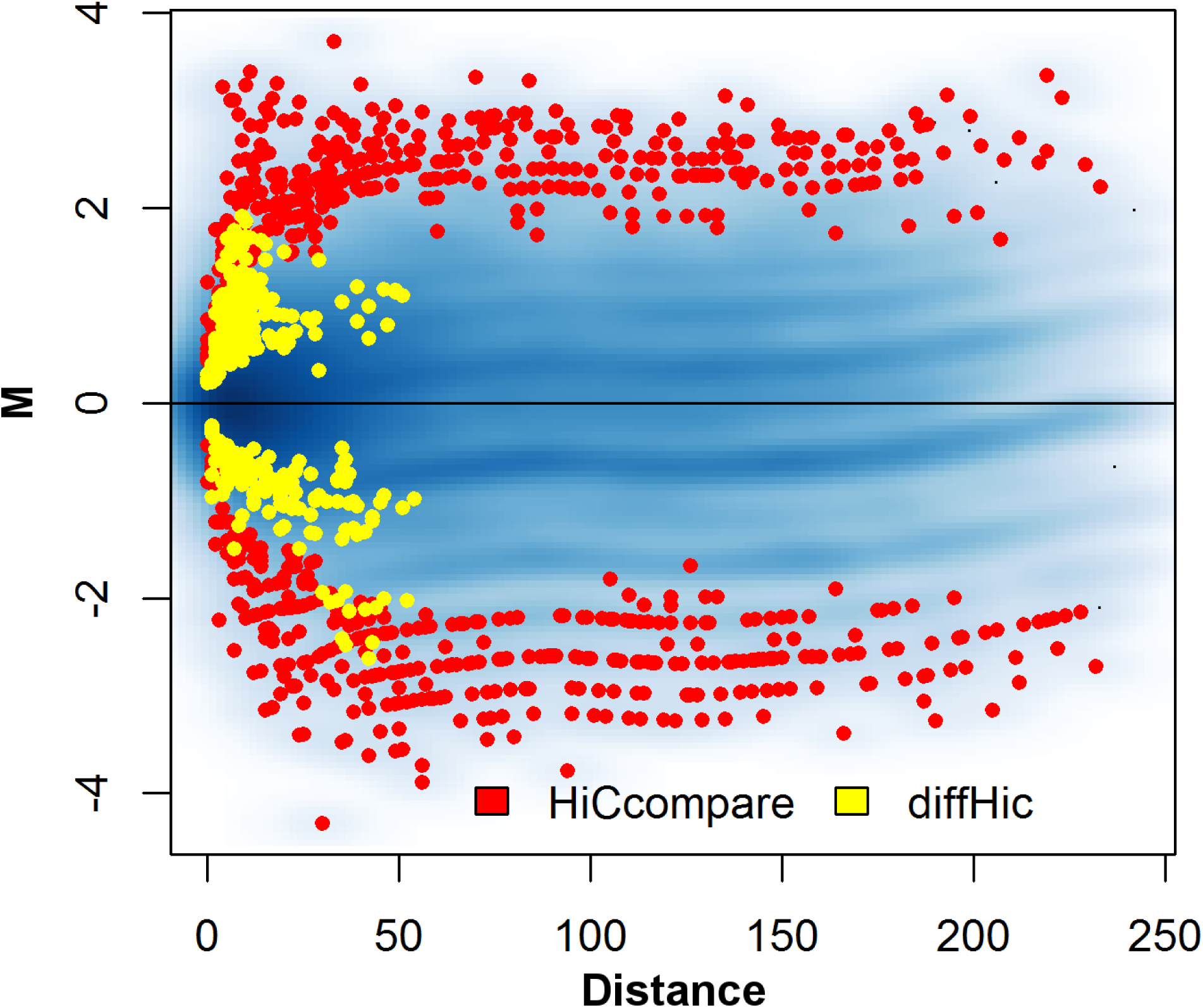
Comparison of regions detected by HiCcompare and diffHic. Chromosome 1 at 1MB resolution for the comparison of RWPE1 prostate epithelial cells and ERG3 over-expression strains of RWPE1 cells. MD plot showing regions detected as significant by `diffHic` in yellow and regions detected as significant by `HiCcompare` in red.

Four pairs of differentially interacting regions were validated using Fluorescence In Situ Hybridization (FISH) [36] and confirmed in the `diffHic` analysis [51]. These regions were also detected as differentially interacting in the `HiCcompare` analysis (Table 1). These results suggest that both methods are able to detect biologically relevant chromatin interaction differences.

**Table 1. Differential interactions validated by FISH detected by HiCcompare and diffHic.** The regions containing genes listed in the “Interaction” column were previously validated by Fluorescence In Situ Hybridization (FISH) as differentially interacting [36]. The “Difference” columns show the differences, measured as *log*2(*IF*_1_*-IF*_2_), detected by the `HiCcompare` and `diffHic` pipelines; the “Significance” columns show the permutation p-value (“HiCcompare”) and FDR-corrected p-value (“diffHiC”) for the corresponding differential interactions.

## Discussion

This work introduces three novel concepts for the joint normalization and differential analysis of Hi-C data, implemented in the `HiCcompare` R package. First, we introduce the representation of the differences between two Hi-C datasets on an MD plot, a modification of the MA plot [41]. Importantly, we consider the data on a per-unit-length-distance basis, allowing normalization of global biases in the CNV-neutral data without distorting the relative distribution of interaction frequencies of the interacting regions. Second, we implement a non-parametric `loess` normalization method that jointly accounts for biases in the normalzied datasets. There is compelling evidence that non-parametric normalization methods, such as quantile- and `loess` normalization, are particularly suitable for removing the between-dataset biases [48,49], confirmed by our application of `loess` to the joint normalization of Hi-C data. Third, we develop and benchmark a simple but rigorous statistical method for the differential analysis of Hi-C datasets.

Our method is designed to analyze processed Hi-C data summarized in a sparse matrix text format (see Methods). There is no *de facto* standard for a compact representation of Hi-C chromatin interaction data, with each major study introducing their own *ad hoc* format [14,39]. A general consensus is to use an extension of the widely used BED format, termed BEDPE (Browser Extensible Data Paired-End) [55]. It contains six mandatory columns corresponding to the chromosome, start and end positions of a pair of regions and, optionally, name, score, strand of both regions. It can be extended with additional columns and easily viewed in any text editor on any platform. A simplified version of this format, PGL, was recently published [56]. Another specialized text format, `.hic`, was designed to store matrices at different resolutions, with an index allowing quick access to any region [39]. The binary representation of Hi-C data was also implemented in the HDF5-based.`cool` (https://github.com/mirnylab/cooler) and `BUTLR` [57] data formats. We designed `HiCcompare` to reformat sparse upper triangular and `.cool`-formatted data into a BEDPE format amenable for straightforward computational processing.

Although `MA` joint normalization uses a similar concept of representing measures from two datasets on a single plot (S4 Fig B), it performed inferior to the `loess` joint normalization. This is likely due to the fact that `MA` normalization represents the data using *A*verage chromatin interaction frequency as an X-axis. However, due to power-law decay of interaction measures, the dynamic range of *A*verage chromatin interaction frequency is limited, making `loess` curve fitting difficult. This limitation leads to differences being detected at short distances (Fig 4, S4 Fig B). Instead, the more balanced representation of chromatin interaction differences *M* (Y-axis) as a function of distance *D* (X-axis) improves the performance of the `loess` fit for the joint normalization and the subsequent detection of chromatin interaction differences.

Although our classification evaluation and ROC curve analyses showed a clear advantage of the `loess` joint normalization, differential chromatin interactions could still be detected in the individually normalized matrices. `KR` and `SCN` were among the top performing individual normalization methods despite the fact they fail to remove biases between datasets (S1 File). Their relatively good performance can be explained by our permutation-based method of detecting differential chromatin interactions. Based on the concept of the MD plot and the per-unit-length-distance permutation testing for differential chromatin interactions, our method is designed to detect differential interactions even in the presence of biases. Still, the superior performance of the `loess` joint normalization indicates the need to jointly account for biases between matrices when detecting differentially interacting chromatin regions.

The current implementation of `HiCcompare` was tested on chromosome-specific chromatin interaction matrices. They are believed to represent the true chromatin interactions arising from distinct chromosome territories [14]. However, a substantial proportion of chromatin interactions (∼10-50%) arise from the inter-chromosomal interactions [13,14,37,39,58]. The extreme variability and poor replicability suggests that such inter-chromosomal contacts result from random collisions among chromosomes. The source and biological relevance of inter-chromosomal interactions remain a topic of intense research [43]. Our future directions include investigating the joint normalization and differential analysis of inter-chromosomal interaction matrices.

Increasing resolution of the size of interacting chromatin regions requires a significant increase in sequencing coverage. To achieve the genome-scale coverage at kilobase-pair resolution conventional Hi-C experiments require billions of DNA sequencing reads [13,39]. Existing Hi-C data at high resolutions still suffer from a limited dynamic range of chromatin interaction frequencies, with the majority of them being small or zero, especially at large distances between interacting regions (S7 File). The problem is exacerbated in single-cell Hi-C technology, which generates very sparse Hi-C data even at 1Mb resolution [59]. This sparsity places limits on `loess` joint normalization, as it builds a rescaling model from many non-zero pairwise comparisons. This sparsity explains our observed sub-optimal performance of `loess` in higher resolution data (S7 File). A way to alleviate this limitation is to consider interactions only within a range of short interaction distances, where genomic regions interact more frequently and the proportion of zero interaction frequencies is the lowest. Decreasing costs of sequencing technologies will eventually overcome the problem of insufficient coverage of high-resolution chromatin interaction matrices, making them amenable for `loess` joint normalization and the detection of differential chromatin interactions.

Despite the ability of Hi-C technology to simultaneously capture all genomic interactions, current resolution of Hi-C data (1Mb - 1kb) remains insufficient to resolve individual *cis*-regulatory elements (∼100b-1kb). Alternative techniques, such as ChiA-PET [60], Capture Hi-C [8] have been designed to identify targeted 3D interactions, e.g., between promoters and distant regions. These data require specialized normalization methods [22]. Our future goals include extending the `loess` joint normalization method for chromosome conformation capture data other than Hi-C.

## Methods

### Data

A chromosome-specific Hi-C chromatin interaction matrix is a square matrix of size *N xN*, where *N* is the number of genomic regions of size *X* on a chromosome. The size *X* of the genomic regions defines the resolution of the Hi-C data. Each cell in the matrix contains an interaction frequency *IF*_*i,j*_, where *i* and *j* are the indices of the interacting regions.

A full chromatin interaction matrix is symmetric around the diagonal, and sparse, i.e., containing many zero interaction frequencies. As such, it can be condensed into a sparse upper triangular matrix without loss of information and with the benefit of a smaller file size. The sparse upper triangular matrix contains three columns: index *i* of the first region in the pair, index *j* of the second region, and the non-zero interaction frequency *IF*_*i,j*_. Functions to convert the full chromatin interaction matrix into a sparse format and back are provided. For this study, data in the sparse upper triangular format from the GM12878, K562, IMR90, HMEC, NHEK, and RWPE1 cell lines were used (S1 Table).

### Visualization of the differences between two Hi-C datasets using an MD plot

The first step of the `HiCcompare` procedure is to convert the data into what we refer to as an MD plot. The MD plot is similar to the MA plot (Bland-Altman plot) commonly used to visualize gene expression differences [41]. In terms of gene expression, the MA plot visualizes gene expression differences (**M**inus) at a given expression level (**A**verage). MA plots have been used to visualize and normalize ChIP-seq data [49,50] and Hi-C data [51] using the same principle of fitting a `loess` curve through the plot and rescaling both datasets. Instead of **A**verage, the MD plot uses distance between interacting regions (**D**istance).

*M* is defined as the log difference between the two data sets *M* = *log*_2_(*IF*_2_*/IF*_1_), where *IF*_1_ and *IF*_2_ are interaction frequencies of the first and the second Hi-C datasets, respectively. *D* is defined as a distance between the interacting regions, expressed in unit-length of the *X* resolution of the Hi-C data. In terms of chromatin interaction matrices, *D* corresponds to the off-diagonal traces of interaction frequencies (Fig 1A). Because chromatin interaction matrices are sparse, i.e. contain an excess of zero interaction frequencies, by default only the non-zero pairwise interaction are used for the construction of the MD plot with an option to use partial interactions, i.e. with a zero value in one of the matrices and a non-zero IF in the other.

### Joint normalization of multiple Hi-C data using loess regression

After the transformation of the data into an MD plot, `loess` regression [47] is performed with *D* as the predictor for *M*. The `loess.as` function from the `fANCOVA` R package is used for the `loess` step. We use a first-degree polynomial regression with the generalized-cross validation (gcv) automatic smoothing parameter selection. The automatic smoothing parameter selection process determines the optimal span for the `loess` regression to be used for the whole dataset. For the Hi-C data tested, typical spans were between 5-12%. Once the `loess` model is fit, we use the predicted values to normalize the original IFs in the chromatin interaction matrices.

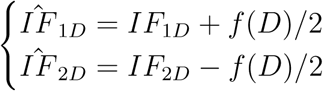

where *f* (*D*) is the predicted value from the `loess` regression at a distance *D*. Note that for both Hi-C datasets the average interaction frequency remains unchanged, as the one set is increased by the factor of *f* (*D*)*/*2 while the other is decreased by the same amount. The normalized matrices are then anti-log transformed.

### Normalization methods for individual Hi-C datasets

Four methods for normalizing individual Hi-C datasets were compared with the `loess` joint normalization method. Here, we briefly describe them.

The `ChromoR` method [9] applies the Haar-Fisz Transform (HFT) to decompose a Hi-C contact map. HFT assumes the IFs in the contact map are distributed as a Poisson random variable. After HFT decomposition, wavelet shrinkage methods for Gaussian noise are applied for de-noising. The contact map is then reconstructed with the inverse HFT. The `ChromoR` R package was used to normalize the matrices with the `correctCIM` function.

ICE (iterative correction and eigenvector decomposition) normalization [30] functions by modeling the expected *IF*_*ij*_ for every pair of regions *(i,j)* as *E*_*ij*_ = *B*_*i*_*B*_*j*_*T*_*ij*_, where *B*_*i*_ and *B*_*j*_ are the biases and *T*_*ij*_ is the true matrix of normalized IFs. The maximum likelihood solution for the biases *B*_*i*_ is obtained by iterative correction. It attempts to make all regions equally visible, and was shown to perform as well as the explicit bias correction method by Yaffe and Tanay [61]. `ICE` normalization was performed using the `HiTC` R package’s `normICE` function.

`KR` (Knight-Ruiz) normalization [31] is another “equal visibility” algorithm that balances a square non-negative matrix *A* by finding a diagonal scaling of *A* such that *P* = *D*_1_*AD*_2_ sums to one. The `KR` algorithm uses an iterative process to find *D*_1_ and *D*_2_ scaling matrices by alternately normalizing columns and rows in a sequence of matrices using an approximation of Newton’s method. The `KR` normalization method was re-implemented in R using the published `matlab` code [31] and is included in the `HiCcompare` package as the `KRnorm` function.

`SCN` (Sequential Component Normalization) [29] is a method that is broadly generalizable to many Hi-C experimental protocols. It attempts to smooth out biases due to GC content and circularization. `SCN` works by first normalizing each column vector of a Hi-C contact matrix to one using the Euclidean norm. Then each row of the resulting matrix is normalized to one using the row Euclidean norm. This process is repeated until convergence (usually 2 to 3 iterations). The SCN method was re-implemented in R and included in the `HiCcompare` package as the `SCN` function.

`MA` (Minus Average normalization) [51] is a commonly used normalization method for genomic data. It is based on the MA plot where the data is plotted according to the Average log counts (or counts per million) and the log Minus (difference) between the two data sets. A `loess` model is then fit to this plot and the residuals for the fit can be used to smooth the data sets. MA normalization was implemented in R and included in the `HiCcompare` packages as the `MA_norm` function.

### Detection of differential chromatin interactions

After joint normalization, the normalized chromatin interaction matrices are ready to be compared for differences. Again, the MD plot is used to represent the differences *M* between two normalized datasets at a distance *D*. Only the non-zero pairwise interaction frequencies are visualized and tested for significant differences. At a given distance *d*, each difference *M*_*id*_ is tested for significance using a permutation test:

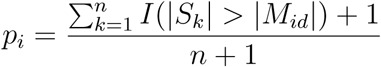

where *S* is a sample of size *n*, taken with replacement, of differences sampled from a vector of *M*_*d*_, and *M*_*id*_ is the *i*^*th*^ difference tested for significance. *I* i s the i dentity f unction. S ince the number of differences diminishes with the increasing genomic distance (less off-diagonal IFs in the upper right corner of interaction matrices), differences for the top 15% of distances are combined to have a pool of variables for permutation purposes. Note that the permutation framework also accounts for multiple testing correction. A user-specified significance threshold α(typically, 0.05) is used to define significant differential chromatin interaction frequencies.

The permutation framework tends to detect at least one significant difference at a given unit-distance if the vector, *M*_*d*_, is large enough. In order to reduce the number of false positives, we provide the option to filter the final p-values *pM* _*id*_ by a user specified or automatically calculated fold change *θ*. This option allows for the user to pre-specify the meaningful difference between the two Hi-C datasets that must be reached in order to call a difference truly significant.

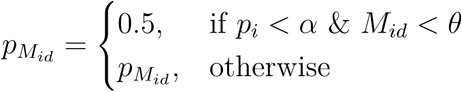

The *θ* threshold is calculated automatically as 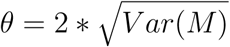, where *M* is the set of all the *M* values from the MD plot. The rationale for a single threshold *θ* is our observation that the standard deviation of the *M* values is approximately constant across the range of distances (S3 Fig). This automatic calculation of *θ* provides a good indicator of the level of noise present between the two datasets and thus any differences detected which fall within the range of (*-θ, θ*) are likely just a result of technical noise and not representative of a truly significant difference between the datasets.

### Estimating power of the differential chromatin interaction detection

The effect of individual vs. joint normalization methods on the power of detection of differential chromatin interactions must be estimated on *a priori* known differences [62]. As there is no “gold standard” for differential chromatin interactions, we created such *a priori* known differences by simulating Hi-C matrices, introducing controlled biases and pre-defined chromatin interaction differences in one of them. The benefit of using joint normalization vs. individually normalized datasets was quantified by the improvement in power of pre-defined chromatin interaction differences using the `pROC` R package. Other standard classifier performance measures (True Positive Rate (TPR), False Discovery Rate (FDR), F1 score, etc.) were also assessed.

#### Simulated Hi-C data

A matrix of chromatin interaction frequencies (*IF s*) at each distance *D* between interacting regions can be represented using four components, 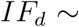 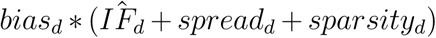. 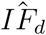 is the expected interaction frequency at a distance *d*, *spread*_*d*_ is the distribution of interaction frequencies at that distance, and *bias*_*d*_ is an optional offset of the interaction frequencies at that distance. Two chromatin interaction matrices of size 100x100 were simulated for the joint normalization and differential chromatin interaction detection.

To model the components used to create simulated chromatin interaction matrices we used the observation that the decay of chromatin interaction frequencies *IF*_*d*_ with increasing distance *d* between interacting regions can be approximated with a power-law distribution *IF* = *C d*^*-α*^ [14,24,26,38,43], where *C* is the constant. The parameter *α* for the first component 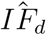 was estimated by fitting the power-law function and optimizing the fit using maximum likelihood estimation. *α* ranged from 1.8 to 2.2 when using datasets from different cutting enzymes (MboI and DpnII) on the cell line GM12878, at resolutions from 1Mb to 50kb, on chromosome 1 (S2 File). The 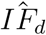 was modeled using *α* = 1.8.

The second component, *spread*_*d*_, represents the distribution of chromatin interaction frequencies (the spread of IFs) at a distance *d*. It was approximated using a normal distribution *N* (0*, SD*), where *SD* is the standard deviation of interaction frequencies *IF*_*d*_ at a given distance *d*. The *SD* parameter was estimated to follow the power-law decay with *α* ranging from 1.6 to 3.2 (S3 File) and set to 1.9 in the current simulations. We found modeling the dependence between SD and the distance between interacting regions is a better approximation then the fixed-step decrease of SD value proposed previously [63]. The 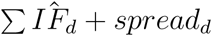 is used to create two matrices with the same underlying signal, but different noise.

Real Hi-C matrices are sparse, that is, they contain zeros. These zero interaction frequencies may arise due to a real lack of interactions, or represent insufficient read coverage or technical artifacts. As such, zero IFs are non-informative and therefore omitted in all calculations. To model the effect of zero IFs, we investigated the dependence of the proportion of zeros vs. distance. Expectedly, the proportions of zeros were minimal at shorter distances between interacting regions, where the probability of interactions is the highest [14]. It increased with the increasing distance, where interactions are less frequent. The higher the resolution of the data (smaller length of the interacting regions) was also found to increase the proportion of zeros. This dependence did not follow a consistent trend other than the fact that the proportion of zeros increases with distance (but may plateau after a point) and with higher resolutions (S4 File). The proportion of zeros was modeled as linearly increasing with distance, *P* (*IF* = 0) = *γ ∗distance*, where the slope, *γ*, is set to 0.001 by default, but can be set by the user in the provided simulation functions. The proportions determined at each unit-length distance were used to set the corresponding number of IFs, sampled uniformly at random, to zero.

The fourth component, *bias*_*d*_, introduces a local offset in one simulated matrix. It is modeled as a systematic deviation of interaction frequencies from the power-law decay. Intuitively, on an MD plot, such a deviation can be represented as a non-linear scaling function across the range of distances between interacting regions. In the current simulation, we modeled bias as a Gaussian function with mean equal to 20 and standard deviation set to 30, creating a “bump” on an MD plot. A user has an option to use any function to model the offset. An optional global scaling can be added to one of the simulated matrices by multiplying all of the IFs by a constant. The global scaling is applied to all interaction frequencies in the matrix, shifting them systematically across the full range of distances. Both components systematically alter the distribution of chromatin interaction frequencies in one of the matrices.

#### Pre-defined chromatin interaction differences

To add known differences to the simulated Hi-C matrices, we introduced known fold changes to one of the matrices. First, the *i, j* coordinates of chromatin interaction frequencies to be changed were defined by taking a random sample with replacement of the *n* matrix row/column indexes. For all simulations we used *n* = 250, resulting in a randomly selected *≤*250 pairwise chromatin contacts. The interaction frequencies at these coordinates were altered as:

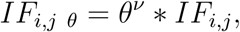

where *θ* is the fold change applied to the cell and

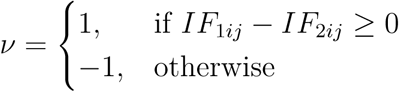

### Excluding CNV regions

Copy number variations (CNVs) can potentially cause issues for normalization and difference detection. To exclude CNV-containing genomic regions a wrapper function was implemented in `HiCcompare` utilizing the `QDNAseq` R package’s CNV detection method [64]. Given two BAM files, this function will check every region of the genome for CNVs and return a call for the CNV status for each region. The regions which are determined to have deletions or amplifications can then be excluded from any further analysis in `HiCcompare` by the user. Alternatively, CNV regions can be detected separately and provided to `HiCcompare` as a BED file. Additionally, the `HiCcompare` package includes the ENCODE blacklisted regions for hg19 and hg38 genome assemblies. These regions, along with CNV regions can be excluded from analyses.

### Comparison with diffHic

A `HiCcompare` analysis was performed on Hi-C data from RWPE1 prostate epithelial cells with differences in ERG3 over-expression [36]. RWPE1-ERG and RWPE1-GFP conditions were compared. Briefly, the sequencing data for library 1 was obtained from GEO (GSE37752 accession number) in the form of FASTQ files. They were aligned to the GRCh37/hg19 version of human genome using the `HiCUP` v0.5.9 pipeline [65]. The resulting BAM files were then converted to HIC files using Juicer tools v0.8 [66]. A CNV detection step was performed using `QDNAseq` v1.12.0 [64]. The HIC files were processed through the `HiCcompare` pipeline at 1Mb resolution as described. Regions with an overlap of at least 20% of the detected CNV regions and/or ENCODE blacklisted regions were excluded from further analysis. `HiCcompare`-detected differentially interacting regions were compared for overlap to regions detected by `diffHic`, provided by Drs. Gordon Smyth and Aaron Lun. The regions detected by `diffHic` were also compared for overlaps with the blacklisted regions and regions detected as CNVs. Lun & Smyth confirmed some of the interactions they detected using FISH. The coordinates for these genes were overlapped with the results of `HiCcompare` and `diffHic` to confirm that they were also detected by each method. Because these genes overlapped with several detected regions in the results of `HiCcompare` and `diffHic`, the interaction with the most overlap of the genes detected by FISH was chosen for the comparison displayed in Table 1.

### Implementation and Availability

`loess` joint normalization and differential chromatin interaction detection methods are freely available as an R package `HiCcompare`, available from Bioconductor https://bioconductor.org/packages/HiCcompare/. Its development continues on the GitHub repository https://github.com/dozmorovlab/HiCcompare. All functions were implemented and tested in R/Bioconductor environment v.3.4.1.

## Acknowledgements

The authors thank Drs. Gordon Smyth and Aaron Lun for providing `diffHic` analysis results. Research reported in this publication was supported in part by the American Cancer Society Institutional Research Grant to MD, and by the National Institute Of Environmental Health Sciences of the National Institutes of Health under Award Number T32ES007334 supporting JS. The content is solely the responsibility of the authors and does not necessarily represent the official views of the National Institutes of Health and the American Cancer Society.

## Supporting Information

**S1 Fig. Deviation from power-law.**

Deviation from an ideal power-law relationship (red line) between the *log*_10_*-log*_10_ interaction frequencies and distance. Chromosome-specific data for Gm12878 cell line, DpnII enzyme, 1MB resolution were used. Each curved line represents chromosome-specific `loess` fit of the relationship. Full range of distances is shown.

**S2 Fig. ROC analysis on real data.**

ROC curves of the differential chromatin interaction detection using different normalization techniques at (A) 1.5, (B) 2.0, (C) 3.0, (D) 4.0 fold changes. Gm12878 chromosome 1, at 1MB resolution with 500 controlled changes added.

**S3 Fig. Approximately constant** *SD* **of the** *M* **differences across** *D* **distances**

Straight lines represent linear fits to the observed *SD*s of *M* across distances *D*. Data from GM12878, K562, NHEK, and IMR90 cell lines from chromosome 1 at 1MB resolution was used for pairwise comparisons.

**S4 Fig. diffHic results in MD and MA plot form**

Chromosome 1 at 1MB resolution for the comparison of RWPE1 prostate epithelial cells and ERG3 over-expression strains of RWPE1 cells. (A) MD plot for the results of `diffHic` with coloring for significance. (B) MA plot for the results of `diffHic`. Log CPM is plotted on the x-axis and the M values are plotted on the y-axis. Coloring based on significance.

**S1 Table. Hi-C data sources.**

**S2 Table. Comparison of `HiCcompare` and `diffHic` results.**

“Chromosome” - results are broken down by chromosome; “Number of differentially interacting region pairs” - counts of region pairs detected as significantly differentially interacting by the “HiCcompare” and “diffHiC” pipelines, “Overlap” between them, counts of “diffHiC regions overlapping CNVs” and “diffHiC regions overlapping blacklisted” regions; “Average positive difference” and “Average negative difference”, measured as *log*2(*IF*_1_*-IF*_2_), detected by the “HiCcompare” and “diffHiC” pipelines.

**S1 File. Normalization method comparison.**

Persistence of bias in individually normalized chromatin interaction matrices, and its effect on the detection of differential chromatin interactions.

**S2 File. Estimation of the IF power-law depencence.**

Estimation of the power-law dependence between the *log*_10_ – *log*_10_ interaction frequencies and the distance between interacting regions.

**S3 File. Estimation of the SD power-law dependence.**

Estimation of the power-law dependence between the *log*_10_*-log*_10_ SD of interaction frequencies and the distance between interacting regions.

**S4 File. Estimation of the proportion of zeros.**

Estimation of the dependence between the proportion of zeros and distance between interacting regions.

**S5 File. Evaluation of difference detection in simulated data.**

Extended evaluation of differential chromatin interaction detection analysis using simulated Hi-C data. “TP” - true positives, “FP” - false positives, “TN” - true negatives, “FN” - false negatives, “True Positive Rate” - aka recall, or sensitivity *T P/*(*T P* + *F N*), “Specificity” - *T N/*(*F P* + *T N*), “Precision” - *T P/*(*T P* + *F P*), “False Positive Rate” - *F P/*(*F P* + *T N*), “False Negative Rate” - *F N/*(*T P* + *F N*), “False omission rate” - *F N/*(*F N* + *T N*), “Negative Predictive Value” - *T N/*(*F N* + *T N*), “F1” - *F*_1_ score 2*T P/*(2*T P* + *F P* + *F N*), “Accuracy” - (*T P* + *T N*)*/*(*T P* + *F P* + *T N* + *F N*), “AUC” - area under ROC curve.

**S6 File. Evaluation of difference detection in real data.**

Extended evaluation of differential chromatin interaction detection analysis using real Hi-C data. “TP” - true positives, “FP” - false positives, “TN” - true negatives, “FN” - false negatives, “True Positive Rate” - aka recall, or sensitivity *T P/*(*T P* + *F N*), “Specificity” - *T N/*(*F P* + *T N*), “Precision” - *T P/*(*T P* + *F P*), “False Positive Rate” - *F P/*(*F P* + *T N*), “False Negative Rate” - *F N/*(*T P* + *F N*), “False omission rate” - *F N/*(*F N* + *T N*), “Negative Predictive Value” - *T N/*(*F N* + *T N*), “F1” - *F*_1_ score 2*T P/*(2*T P* + *F P* + *F N*), “Accuracy” - (*T P* + *T N*)*/*(*T P* + *F P* + *T N* + *F N*), “AUC” - area under ROC curve.

**S7 File. `loess` at varying resolution.**

Visualization of the `loess` joint normalization over varying resolutions.

